# mRNA vaccination overcomes hemozoin-mediated impairment of whole parasite vaccine efficacy for malaria

**DOI:** 10.1101/2025.10.29.685344

**Authors:** Mariah Hassert, Lisa L. Drewry, Lecia L. Pewe, Lisa S. Hancox, Rui He, Sahaana Arumugam, Madison R. Mix, Aliasger K. Salem, John T. Harty

## Abstract

Malaria is a persistent global public health threat. Immunization with Radiation Attenuated Sporozoites (RAS) drives efficient CD8+ T cell dependent sterilizing immunity to malaria in humans. However, this protection is compromised in individuals living in malaria endemic regions and the mechanism(s) of the vaccine failure remain incompletely understood. In this study, we develop a murine model of *Plasmodium* infection where prior blood-stage exposure compromises RAS-induced CD8+ T cell responses and subsequent protection. We further identify the persisting malarial pigment hemozoin as a mediator of impaired CD8+ T cell responses. Mechanistically, we link this defect to impaired antigen uptake by dendritic cells, leading to reduced T cell activation. Importantly, we designed a lipid nanoparticle-encapsulated mRNA vaccine that encodes a string of *Plasmodium* CD8+ T cell epitopes and found that this vaccine overcomes the T cell defect and restores protection in *Plasmodium* exposed mice. Moreover, a combined RAS+mRNA vaccine regimen enhances liver resident memory T cells and protection even in malaria experienced hosts. These findings support the identification of hemozoin as a long-lived obstacle to vaccine efficacy in malaria endemic areas and provide a rational framework for designing malaria vaccines that are effective in endemic settings.

## INTRODUCTION

Mosquito-borne *Plasmodium* infections remain a global health crisis, with >200,000,000 cases and >600,000 fatalities per year^1^. *Plasmodium* infections are initiated by infected anopheline mosquitoes during a blood meal, where the parasites are deposited in dermal tissues. Parasites then invade the vasculature and rapidly move to the obligatory, yet asymptomatic hepatocyte infections of the liver-stage before transitioning to infection of red blood cells during the symptomatic blood-stage of infection. Development of a highly efficacious vaccine targeting the asymptomatic liver-stage of malaria would greatly reduce disease burden by eliminating the morbidity, mortality, and transmission that are caused by the subsequent blood-stage of malaria^2^.

<u>R</u>adiation <u>A</u>ttenuated <u>S</u>porozoite vaccination (RAS) has been studied for decades and categorically termed the “Gold Standard” liver-stage malaria vaccine^3^. Attenuation of sporozoites through radiation generates immunity against liver-stage *Plasmodium* infection by triggering an abortive, but immunogenic infection— terminating prior to the initiation of blood-stage^4,5^. In murine, nonhuman primate and human challenge models, RAS has shown profound efficacy; being capable of delivering up to 100% sterilizing immunity^6–9^. RAS immunization elicits both circulating (T_CIRCM_) and liver resident (T_RM_) memory CD8+ T cells which have been determined by multiple studies to be the major mechanism of protection induced by this vaccine^10–17^.

Several malaria vaccine candidates, including RAS, work well in malaria-naïve humans, but show between 30-50% efficacy when tested in field trials in endemic regions^18,19^. A number of non-mutually exclusive hypotheses have emerged to explain this phenomenon including antigenic variability in field parasites, differences in environmental factors, host genetic factors, and the history of antigen exposures^20–22^. One additional hypothesis posits that prior malaria exposure may negatively affect the immune responses of people living in endemic areas, though the exact mechanisms remain unknown^23^. Here, we further developed a mouse model^24^ showing prior blood-stage *Plasmodium* exposure compromised RAS-induced CD8+ T cell responses, and ultimately compromised vaccine-mediated protection from subsequent challenge infection. This long-lasting immune perturbation persisted for more than one year and was conserved after blood-stage infection with different *Plasmodium* species suggesting common mechanistic underpinnings. One conserved feature of *Plasmodium* blood-stage infections is the formation of hemozoin, a persisting biocrystal produced to detoxify heme during the digestion of hemoglobin in infected red blood cells^25^. Native hemozoin purified from infected red blood cells has immunostimulatory capacity that depends on contamination with parasite DNA and signaling through TLR9 or NLRP3^26–28^. To address the role of persisting hemozoin in the absence of contaminants, we injected mice with synthetic hemozoin then studied their T cell responses and protection after RAS vaccination. Injection of mice with synthetic hemozoin alone phenocopied prior infection, implicating this long-lasting parasite-produced metabolic byproduct in compromised CD8+ T cell responses and poor vaccine efficacy. We found that the blood-stage-induced T cell defect could be rectified by multiple strategies, including LNP-mRNA vaccination. Moreover, combining translationally relevant RAS and LNP-mRNA vaccines enhanced protective *Plasmodium*-specific T resident memory T_RM_ formation in the liver. This newfound understanding of the role of hemozoin in modulation of T cell priming during liver-stage vaccination identifies a critical obstacle and highlights potential strategies for anti-malarial vaccines with efficacy in malaria endemic regions.

## RESULTS

### Reduction in RAS-induced CD8+ T cell responses drives impaired vaccine efficacy in Plasmodium experienced mice

To model the impact of prior malaria exposure on immune responses to RAS, we injected mice with infectious sporozoites (not shown) or infected red blood cells (iRBC) of the non-lethal parasite strain *P. yoelii* 17XNL (*Py*), which is cleared from C67BL/6J mice in 25-30 days^29^. After parasite clearance, mice were vaccinated with RAS from a different *Plasmodium* species (*P. berghei ANKA*) expressing the model antigen ovalbumin (*Pb*-Ova^30^) to elicit transgenic OT-I^31^ or endogenous RPL6 epitope^11^-specific CD8+ T cell responses (**Fig. 1A**, **Extended Data Fig.1**). We observed a 2-10-fold numerical reduction in circulating effector CD8+ T cell responses (**Fig. 1B, 1C**) as well as splenic (T_CIRCM_) and liver memory T cells responses (**Fig. 1D, 1E**) in *Py* experienced mice from both sporozoite- (not shown) and blood-stage infections. This indicates that the immune compromise was caused by blood-stage infection. Of note, the OT-I sensors evaluated here were injected after clearance of the parasite and were not exposed to malaria, showing that the impairment in their response appears to be T cell extrinsic. Similar *Pb*-Ova RAS parasite burdens were observed in livers of *Py*-naïve and experienced mice, so a reduction in liver-stage infection/antigen load through cross-species immunity is unlikely to account for these outcomes (**Extended Data Fig. 2**). Upon challenge with virulent *Pb* sporozoites expressing luciferase (*Pb*-Luc), RAS immunized *Py* naïve mice exhibited the expected reduction in liver parasite burden compared to non-immune controls (**Fig. 1F**). However, RAS immunization of *Py* experienced mice did not engender significant immunity (**Fig. 1F**). Consistent with this result, we observed a significant reduction in blood-stage parasite burden at day 5 post challenge^32^ in RAS vaccinated *Py* naïve mice, which was less robust in the *Py* exposed group (**Fig. 1G**). Of note, humans are subjected to multiple booster immunizations with RAS^19^. In general, RAS prime and boost elicits sterilizing immunity against *Pb* in B6 mice^33^. Indeed, RAS boosting further reduced parasite burden in *Py* naïve mice but did not achieve the same efficacy in *Py* experienced mice (**Extended Data Fig. 3**). These findings indicated that prior blood-stage malaria exposure reduced the magnitude of the CD8+ T cell response and vaccine efficacy after RAS immunization— thus providing a model to evaluate poor vaccine efficacy for humans in malaria endemic regions.

**Fig. 1:**
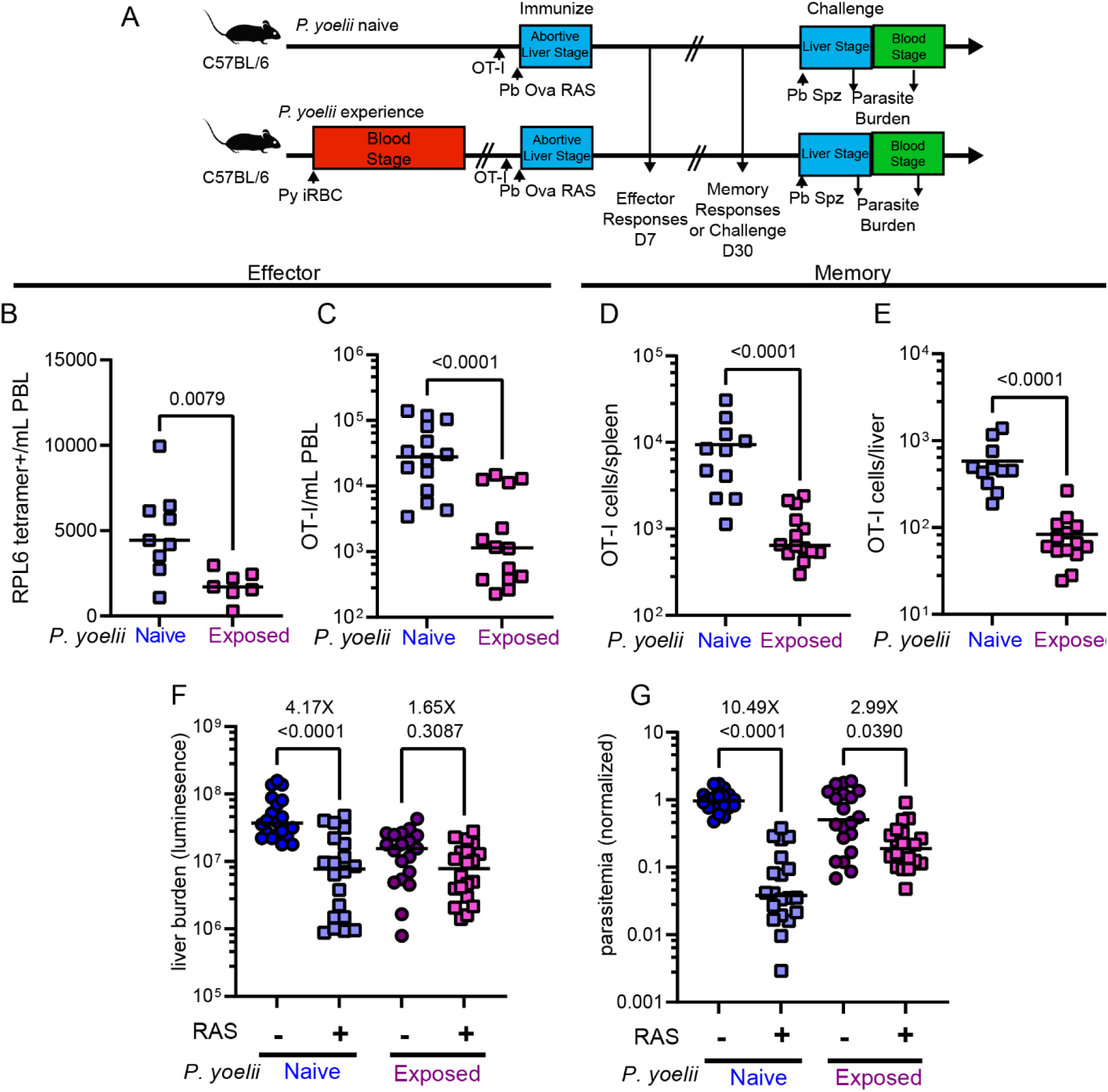
Reduction in RAS induced CD8+ T cell responses drives impaired vaccine efficacy in Plasmodium experienced mice. (A) Experimental design. C57BL/6 mice were infected with 10^6^ *P. yoelii* iRBCs. After blood-stage clearance (30 days), mice were administered 10,000 OT-I cells and vaccinated with 10^4^ *P. berghei* ANKA ova mCherry RAS. 7 days after RAS immunization, blood was collected to evaluate circulating effector CD8+ T cell responses to vaccination. 30 days after blood-stage RAS immunization, spleens and livers were harvested to evaluate memory CD8+ T cell responses. A subset of mice were challenged with 10^4^ virulent *P. berghei*-Luc. (B) Quantification of effector RPL6 specific CD8+ T cells in PBL identified by tetramer staining. (C) Quantification of effector OT-I-tg cells identified by CD90.1 staining. (D) Quantification of memory OT-I cells in the spleen 30 days post RAS immunization. (E) Quantification of memory OT-I cells in the liver 30 days post RAS immunization. (F) Liver parasite burden 40-44 hours after virulent *P. berghei* challenge measured by bioluminescence imaging of whole animals and represented as total flux (p/s). Fold change is listed above p value for each comparison. (G) Relative parasitemia 5 days post virulent *P. berghei* challenge. iRBCs per ml of blood were normalized to the *P. yoelii naïve*, non-immunized control mice. Fold change is listed above p value for each comparison. Data are from 2-4 individual experiments with 7-20 mice per group as indicated by individual symbols.

### Reduction in RAS-induced CD8+ T cell responses is reflected in liver T_EM_ and T_RM_ populations

Both T_EM_ and liver T_RM_ have been demonstrated to be critical for RAS-mediated protection^10,12^. We further subdivided the total memory OT-I cells isolated from the liver into T_CM_, T_EM_, and T_RM_ phenotypes based on CD69, CXCR6, and CD62L expression and found the significant numerical reduction in *Py* experienced mice was predominantly accounted for by the T_EM_ and T_RM_ populations and not the T_CM_ population (**Fig. 2A-D**). Importantly, IFNγ production has also been identified as a correlate of protection from liver-stage malaria^8^. Therefore, we isolated liver lymphocytes 30 days after immunization and evaluated IFNγ production following ex vivo Ova peptide restimulation. We found a significant decrease in the number of IFNγ producing OT-I cells in the livers of *Py* experienced mice (**Fig. 2E**). To identify the cells that accounted for this decrease, we subsetted these cells into T_CM_, T_EM_, and T_RM_. There was a significant decrease in IFNγ producing OT-I T_EM_ and a trending decrease in the number of IFNγ producing OT-I T_RM_ in *Py* experienced mice, but no difference in the number of IFNγ producing T_CM_. (**Fig. 2F-H**). These data support the notion that prior blood stage malaria exposure impairs the generation of protective CD8+ T cell responses to RAS immunization.

**Fig. 2:**
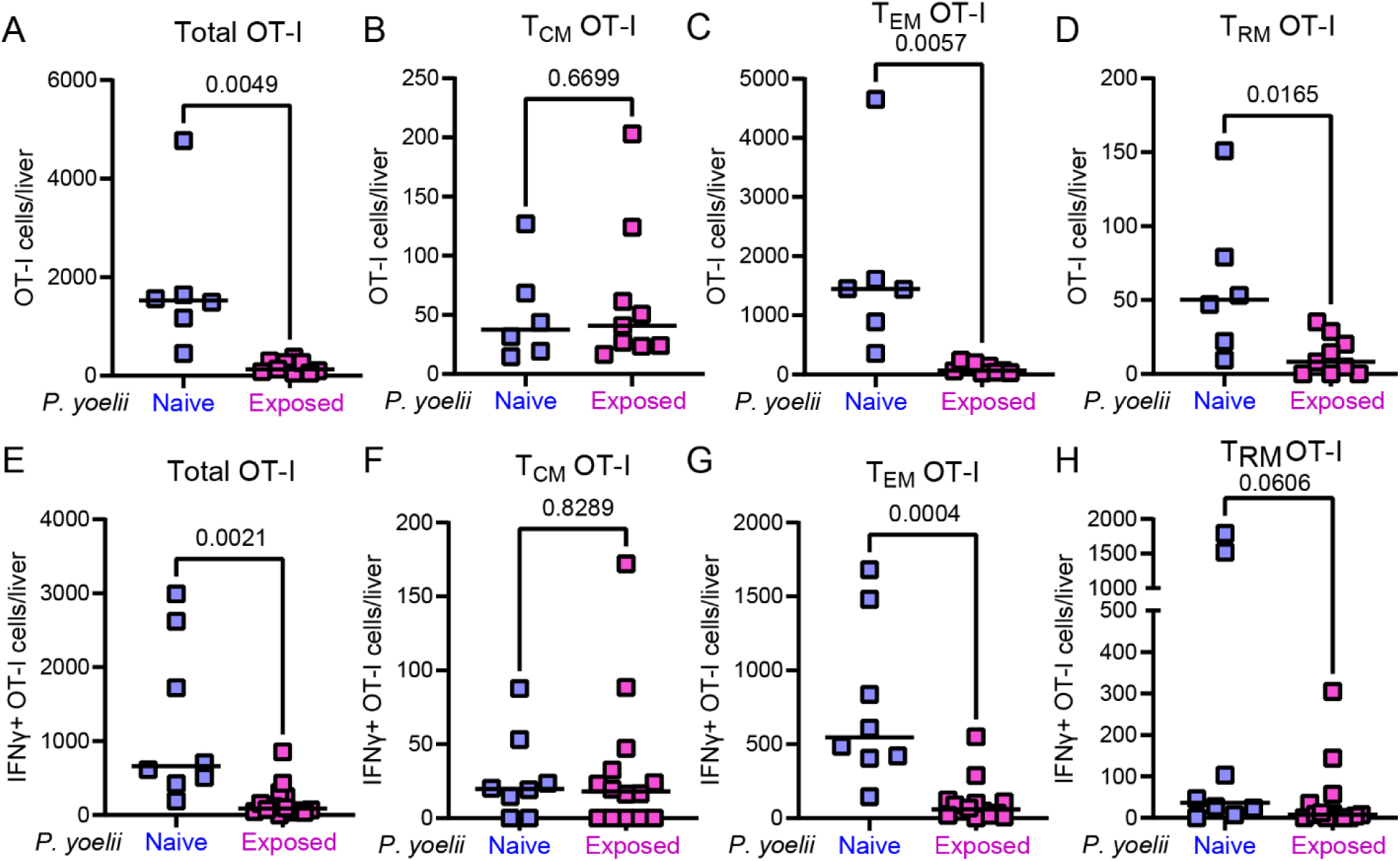
Reduction in RAS-induced CD8+ T cell responses is reflected in liver T_EM_ and T_RM_ populations. Liver memory T cell phenotyping. 30 days post immunization, livers were harvested from mice. (**A**) Total OT-I cells isolated from livers. (**B**) Total T_CM_ OT-I (Lymph→Live→CD8+→ CD90.1+→CD62L+) (**C**) Total T_EM_ OT-I (Lymph→Live→CD8+→ CD90.1+→CD62L-, CD69-, CXCR6-). (**D**)Total T_RM_ OT-I (Lymph→Live→CD8+→ CD90.1+→CD62L-, CD69+, CXCR6+). (**E-H**) Liver lymphocytes were stimulated for 6 hours with ova peptide in the presence of brefeldin A prior to fixation and staining for IFNγ. Data are from 2 independent experiments. (n=6-13 mice per group).

### Impaired RAS immunogenicity and efficacy in blood-stage experienced mice is conserved across Plasmodium strains

The reason(s) why prior *Plasmodium* exposure compromises new CD8+ T cell responses to RAS immunization remain obscure. To gain insight, we asked if the impaired immune responses were specific to *Py* infection or generalizable to other *Plasmodium* species. Injection of *P. chabaudi* chabaudi (*Pcc*) infected red blood cells causes a blood-stage infection lasting ∼15 days with subsequent periodic recrudescence of parasitemia that can be eliminated by antimalarial drug treatment^34^. We then determined whether persistent or chloroquine cured *Pcc* blood-stage infection would also compromise T cell responses and protection after *Pb*-Ova RAS immunization (**Fig. 3A**). Similar to *Py* exposed mice, mice with prior *Pcc* blood stage infection, with or without chloroquine treatment had impaired *Pb*-Ova RAS induced effector CD8+ T cell responses (**Fig. 3B**) and exhibited reduced control of *Pb* sporozoite challenge. (**Fig. 3C, 3D**). These findings indicated that malaria-associated impairments of CD8+ T cell responses to RAS vaccination and impaired vaccine efficacy were conserved after blood-stage infection across rodent *Plasmodium* species.

**Fig. 3:**
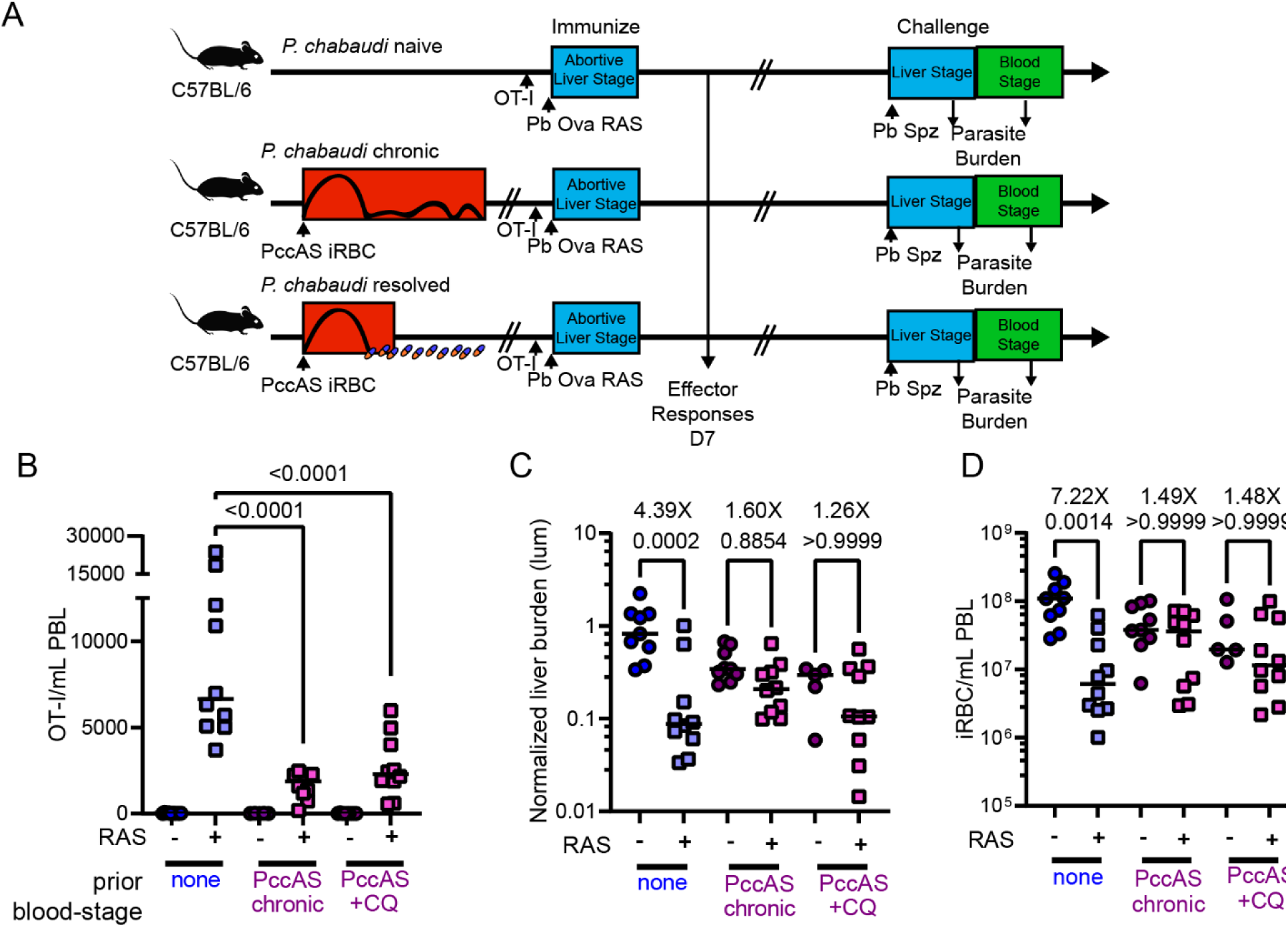
Impaired RAS immunogenicity and efficacy in blood-stage experienced mice is conserved across Plasmodium strains. (A) Experimental design. C57BL/6J mice were infected with 10^6^ *P. chabaudi* AS-mCherry-infected iRBCs Chronic *P. chabaudi* AS-mCherry infection was cured in a subset of mice by administering 600 microgram chloroquine 10x. 60 days after CQ treatment, mice were administered 10,000 OT-I cells and RAS vaccinated. 30 days after RAS vaccination, mice were challenged with 10^4^ virulent *P. berghei* luc sporozoites. (B) Quantification of effector OT-I cells identified by CD90.1 staining of PBL at 7 days post vaccination. (C) Relative liver parasite burden 40-44 hours after virulent *P. berghei* challenge was measured by IVIS. Fold change is listed above p value for each comparison. (D) Breakthrough parasitemia was measured 5 days post virulent *P. berghei* challenge. Fold change is listed above p value for each comparison. Data are pooled from 2-3 independent experiments with 5-13 mice per group as indicated by individual symbols.

### Hemozoin is a candidate driver of impaired RAS efficacy

Controlled human malaria challenge studies (CHMI) indicate that impaired immunity in malaria experienced individuals may be derived from parasite factor(s) capable of modulating the immune response^35,36^. To gain further insight, we infected some mice with *Py* and then immunized these mice and age matched controls 13-19 months later with *Pb*-Ova RAS. *Py* experienced mice still had reduced effector OT-I CD8+ T cell responses following RAS vaccination relative to age-matched controls at these late time points (**Fig. 4A**). Thus, the parasite factor(s) that modulate CD8+ T cell responses to RAS persist long-term after *Py* infection. To further identify characteristics of these factors, we injected fixed iRBC’s from *Py* infected mice or RBCs from influenza strain A/PR/8 infected mice and measured T cell responses to *Pb* RAS-Ova. We observed impaired CD8+ T cell responses to RAS in the mice which received fixed iRBCs from *Py* infected mice even though the mice did not experience an actively replicating blood-stage infection (**Extended Data Fig. 4**). Thus, a factor contained within *Plasmodium* infected RBC was responsible for impaired CD8+ T cell responses in malaria exposed hosts.

**Fig. 4:**
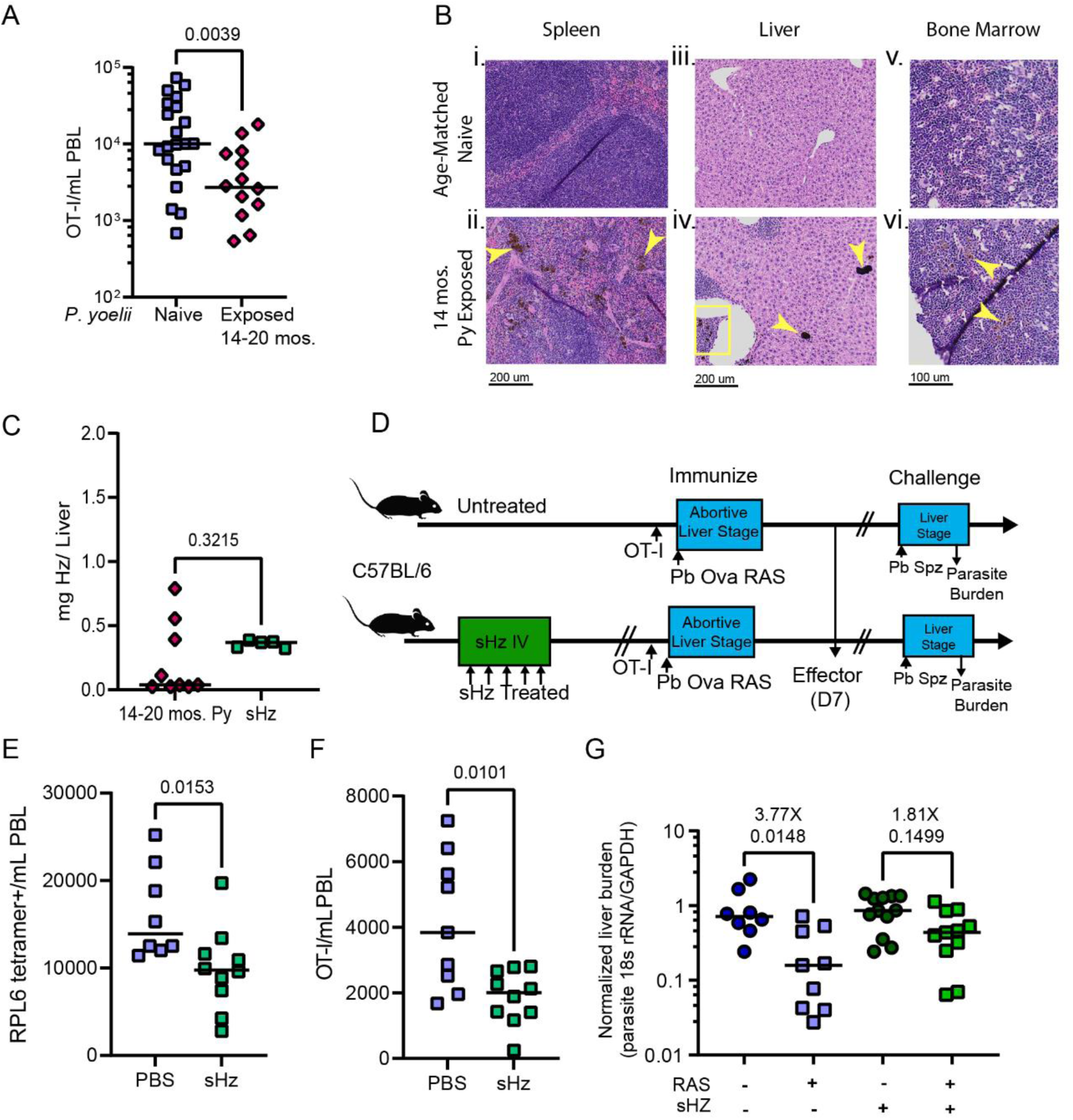
Hemozoin is a candidate driver of impaired RAS efficacy. (A) Quantification of effector OT-I 7 days post RAS immunization of *P. yoelii* experienced mice 14-19 months after parasite clearance or age-matched controls. (B) Representative H&E images of (i) age-matched control spleen, (ii) *P. yoelii* experienced spleen 16 months after parasite clearance, (iii) age-matched control liver, (iv) *P. yoelii* experienced liver 16 months after parasite clearance, (v) age-matched control bone marrow, and (vi) *P. yoelii* experienced bone marrow 16 months after parasite clearance. Examples of Hz deposits are indicated with yellow arrows and boxes. (C) Quantification of Hz present in the livers of mice 14-20 months *post P. yoelii infection*, or 30 days after the conclusion of sHz treatment. (D) C57BL/6 mice were injected IV with 1 mg of sHz five times over the course of 1 week, totaling 5 mg of sHz administered per mouse. 30 days after the last sHz dose, mice were administered 10,000 OT-I cells and were vaccinated with 10^4^ *P. berghei* ANKA ova mCherry RAS. 7 days after RAS immunization, blood was collected to evaluate circulating effector CD8+ T cell responses to vaccination. 30 days after RAS immunization, mice were challenged with 10^4^ virulent *P. berghei*-Luc. (E) Quantification of effector RPL6-specific cells in the PBL 7 days post RAS immunization of sHz treated mice compared to vehicle control. (F) Quantification of effector OT-I cells in PBL 7 days post RAS immunization of sHz treated mice compared to vehicle control. (G) Relative liver parasite burden 42 hours after virulent *P. berghei* challenge was measured by qRT-PCR. Data are from 2 independent experiments. n=4 mice per group for histology. n= 8-21 mice per group for immunological readouts as indicated by individual symbols. n=5-9 mice per group for Hz quantification as indicated by individual symbols.

Together, our results show that the parasite factor(s) that modulate CD8+ T cell responses to RAS are 1) conserved between parasite species, 2) persist for extended periods and 3) are found within infected RBCs. This information led us to investigate a role for the malarial pigment hemozoin (Hz) in the defective CD8+ T cell response. During blood-stage malarial infection, *Plasmodium* parasites catabolize hemoglobin in iRBCs, generating toxic heme moieties^37^. To detoxify heme, the parasite catalyzes formation of the insoluble brown hemozoin (Hz) pigment, which can be detected in infected red blood cells during active infection^38,39^ and becomes localized to phagocytes and extracellular spaces in the liver, spleen and bone marrow after clearance of infection^40^. Hz production is conserved across rodent and human malarial species and has been documented to be long-lived; being detectable in tissues for at least 9 months after infection^41,40^. To extend these data, we observed the persistence of Hz in the spleens, livers, and bone marrow of the mice at 14 months post infection (**Fig. 4B**), correlating with the duration of the observed CD8+ T cell defect in response to RAS priming.

Prior studies have implicated native Hz (nHz) purified from iRBCs as an innate immune modulator, signaling potentially through NLRP3 inflammasome activation^27,42^ and TLR9^43^. However, those observations are complicated to interpret, as nHz is contaminated with parasite nucleic acids or proteins which can have their own discrete bioactivity^26,44^. Indeed, the innate *stimulating* capacity of nHz is eliminated by DNase digestion^45^. However, given that we observed sustained immune impairment out to 19 months post *Py* clearance, it is possible that Hz possesses some discrete immunomodulatory capacity that does not depend on parasite-derived contaminants. To assess the role of Hz, we determined that 5 mg of chemically synthesized hemozoin (sHz) injected intravenously yielded sHz deposition in the liver that was similar to the concentrations found in mice 14-20 months post *Py* exposure (**Fig. 4C**). Consistent with the hypothesis that Hz persistence contributes to impaired CD8+ T cell responses, mice exposed to sHz (**Fig. 4D**) had a significant reduction in *Pb*-Ova RAS-induced endogenous RPL6 specific CD8+ T cells (**Fig. 4E**) and transgenic OT-I cells (**Fig. 4F**) and exhibited reduced protection against *Pb*-Luc sporozoite challenge (**Fig. 4G**). Of note, we determined that when measured by qRT-PCR, parasite burden between naïve and sHz experienced mice that were not RAS immunized was equivalent (**Fig. 4G**). In contrast, when luciferase expression was measured by IVIS as a surrogate indicator of parasite levels, we noted that the signal detected in the sHz exposed group was significantly less than the naïve group (**Extended Data Fig. 5**). These differences are likely due to the dark Hz pigment quenching the visible light signal^46^. In total, these data identified Hz as a major parasite induced factor leading to impaired CD8+ T cell responses and impaired vaccine efficacy following RAS immunization.

### mRNA vaccination and RAS act in concert to enhance liver-stage immunity even in blood-stage experienced mice

Currently, there are not treatments to eliminate Hz from humans that have experienced malaria. Thus, Hz persistence is a major barrier to whole parasite vaccine success in malaria endemic regions. An immunization strategy capable of bypassing this Hz mediated immune impairment would be an important target for malaria vaccine design. mRNA based vaccines have been widely applied to humans during the recent HCoV-2 pandemic^47,48^ and a recent study by Ganely et al. described the use of an RPL6 whole protein encoding, adjuvanted lipoplex mRNA vaccine to achieve protection in *Plasmodium* experienced mice^24^. We recently described an LNP mRNA vaccine designed to elicit robust influenza virus specific CD8+ T cell responses by supporting enhanced proteasomal degradation using a non-cleavable mutant ubiquitin and flanking optimal CD8+ T cell epitopes with optimized proteasomal cleavage residues^49^. To build upon these studies, we re-designed this vaccine to encode a string of H2^b^ and H2^d^ specific CD8+ T cell epitopes from *Pb* (S20, Trap130, CS252, RPL6 proteins, and the model antigen Ova), termed “Ub *Pb* Ova,”) (**Fig. 5A**). This construct allowed us to test the hypothesis that mRNA-LNP vaccination could overcome the diminished CD8+ T cell response in *Plasmodium* experienced hosts (**Fig. 5B**).

**Fig. 5:**
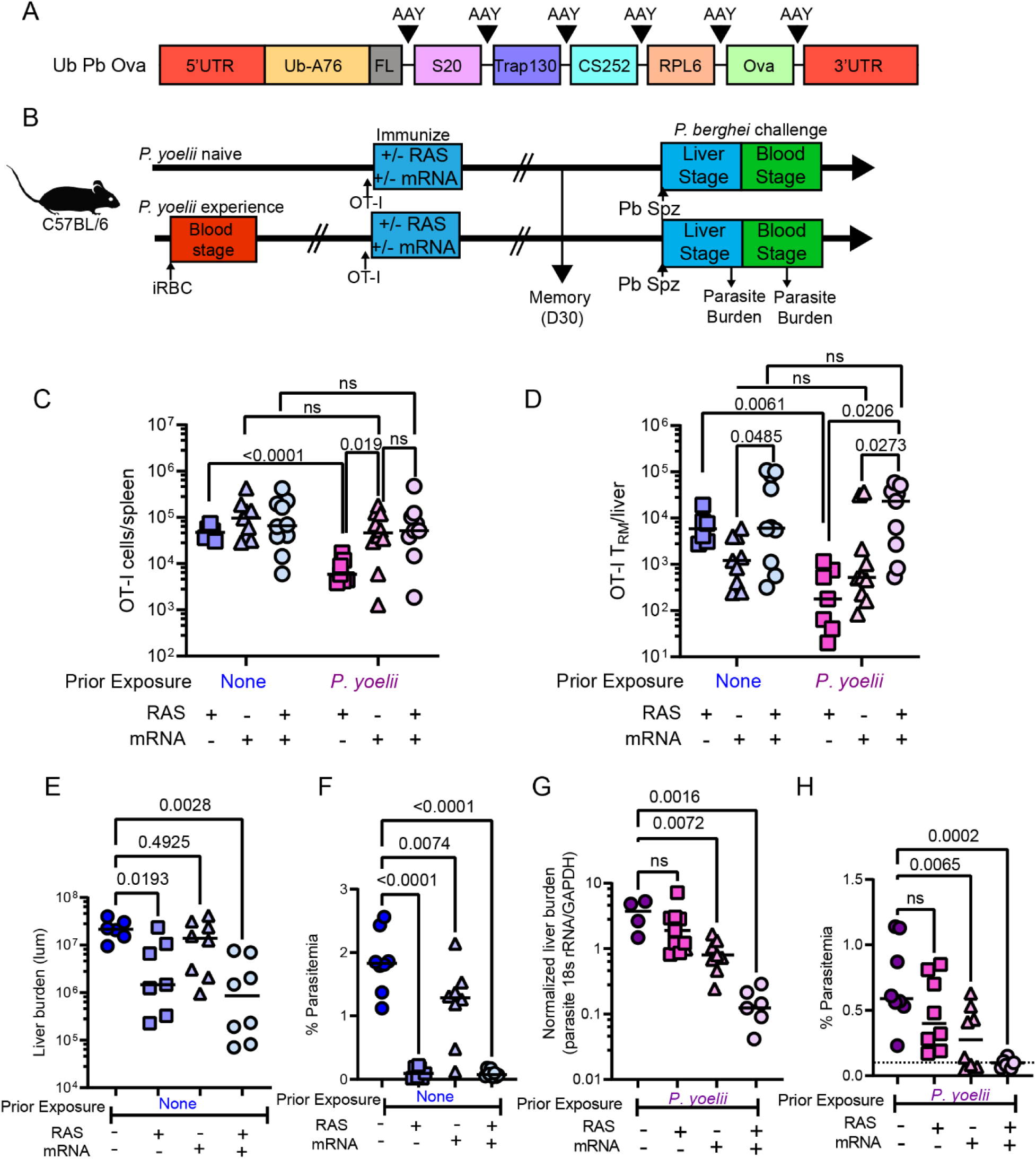
mRNA vaccination and RAS act in concert to enhance liver-stage immunity even in blood-stage experienced mice. (A) Construct design for Ub-Pb-Ova mRNA vaccine. Following the 5’ UTR, a mutant ubiquitin is encoded which cannot be cleaved. Immediately 3’ to the ubiquitin is a flexible linker, followed by the coding sequence for S_20_, Trap_130_, CS_252_, RPL6, and Ova optimal epitopes all flanked by AAY optimal proteasomal cleavage sites. (B) Experimental design. C57BL/6 mice were infected with 10^6^ *P. yoelii* iRBCs or left unmanipulated. After blood-stage clearance (30 days), mice were administered 10,000 OT-I cells and vaccinated with 10^4^ *P. berghei* ANKA ova mCherry RAS, 5 μg of Ub Pb Ova, or both. 30 days after RAS immunization, spleens and livers were harvested to evaluate memory CD8+ T cell responses. A subset of mice were challenged with 10^4^ virulent *P. berghei*-Luc and parasite burden measured by qRT-PCR and breakthrough parasitemia. (C) Quantification of memory OT-I cells in the spleen 30 days post RAS immunization. (D) Quantification of OT-I liver T_RM_ 30 days post immunization. (E) Liver parasite burden in *Py* naïve mice 40-44 hours after virulent *P. berghei-*Luc challenge via IVIS. (F) Percent parasitemia 5 days post virulent *P. berghei* challenge in *Py* naïve mice. (G) Relative liver parasite burden in *Py* experienced mice 40-44 hours after virulent *P. berghei* challenge (parasite 18s RNA/host GAPDH) measured by qRT-PCR. (H) Percent parasitemia 5 days post virulent *P. berghei* challenge in *Py* experienced mice. Data are compiled from 3 independent experiments (n=6-10 mice per group as indicated by individual symbols).

As expected, RAS immunization elicited poor CD8+ T cell responses and reduced protection in *Py* experienced mice compared to *Py* naïve mice (**Fig. 5C-H**). In contrast, mRNA vaccination generated an equivalent memory T cell response in *Py* naïve and experienced mice (**Fig. 5C, D**). Consistent with this, we observed enhanced control of liver-stage *Pb* and delayed breakthrough parasitemia upon virulent sporozoite challenge of the mRNA vaccinated *Py* exposed mice (**Fig. 5G,H**). This indicated that LNP mRNA vaccination of *Plasmodium* experienced mice could overcome the Hz mediated CD8+ T cell impairment. To determine if we could enhance protection by mRNA vaccines, we took advantage of the literature showing that “prime-trap” vaccine strategies utilizing a DNA vaccine prime and RAS boost are capable of enhancing liver T_RM_ formation in naïve mice, which (in addition to T_circM_) are critical in protection against liver-stage malaria^10,12,50,51^. Of note, both mRNA and RAS vaccines are translationally relevant vaccines for humans. Consistent with the literature^50^, combining mRNA vaccination with RAS did not lead to significant increases in circulating memory OT-I responses relative to mRNA alone (**Fig. 5C**). However, combined RAS+mRNA vaccination yielded a significant increase in vaccine specific liver T_RM_ compared to either RAS or mRNA vaccination alone (**Fig. 5D**). Importantly, this dual vaccine administration strategy resulted in equivalent liver T_RM_ responses in naïve vs. *Py* exposed hosts; again demonstrating that the T cell defects associated with Hz exposure could be overcome using LNP mRNA vaccines (**Fig. 5D**). Of note, mRNA vaccination paired with RAS did not impair RAS-induced Ova IgG responses (**Extended Data Fig. 6**). Ultimately, this dual administration vaccine regimen resulted in significantly reduced liver parasite burden (**Fig. 5E and 5G**) and reduced parasitemia (**Fig. 5F and 5H**) in naïve and *Py* exposed mice upon *Pb* sporozoite challenge. We obtained similar outcomes in mice treated with sHz prior to RAS vaccination and challenge (**Extended Data Fig. 7**). These results indicated that not only can mRNA LNP vaccination overcome Hz induced T cell response deficits, but combining mRNA LNP with a single RAS immunization enhanced liver T_RM_ and liver-stage malaria immunity.

### mRNA vaccination bypasses Hz induced antigen uptake defects

The mechanism by which mRNA vaccination can overcome Hz induced vaccine failure is of paramount interest. Our finding that OT-I CD8+ T cells transferred *after* the resolution of blood-stage malaria expand as poorly as endogenous malaria-specific CD8+ T cells suggests that defects in antigen presentation may drive the failure of RAS immunization to efficiently prime CD8+ T cell responses and protect mice against liver-stage malaria. Our lab has previously demonstrated that uptake of hepatocyte-derived *Plasmodium* antigens by CD11c+ APCs in the liver is critical for RAS mediated CD8+ T cell priming^52^. To determine if non-Hz exposed APCs could effectively prime a T cell response in *Py* experienced mice, we immunized *Py* naïve and *Py* experienced mice with activated dendritic cells (DCs) pulsed with Ova peptide (**Fig. 6A**). We found no defect in the elicitation of OT-I effector responses in *Py* experienced mice after DC-peptide priming (**Fig. 6B**), indicating that if functional APCs are present in these mice, correct CD8+ T cell priming can indeed occur. Of note, the CD8+ T cell defect in response to RAS immunization was observed even in *Py* exposed NLRP3-/- mice, suggesting a potentially unique mode of action for Hz that is independent of this innate immune activation pathway (**Extended Data Fig. 8**).

**Fig. 6:**
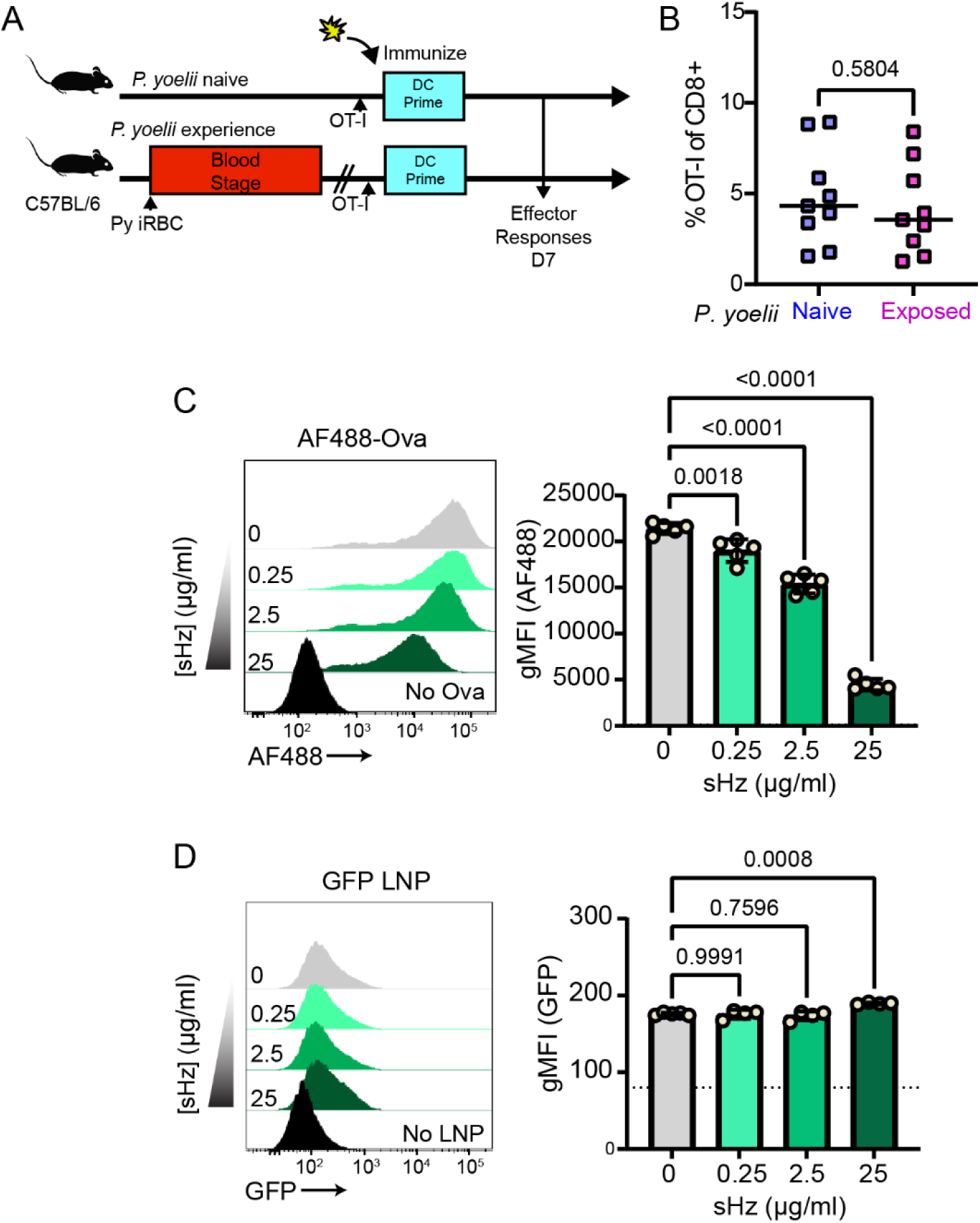
mRNA vaccination bypasses Hz induced antigen uptake defects. (A) DC Prime experimental design. C57BL/6 mice were infected with 10^6^ *P. yoelii* iRBCs. After blood-stage clearance (30 days), mice were administered 10,000 OT-I cells and vaccinated intravenously with 5×10^5^ Ova peptide pulsed activated DCs. 7 days post immunization, blood was collected to evaluate circulating effector CD8+ T cell responses to vaccination. (B) Quantification of effector OT-I cells in PBL 7 days post DC immunization. Data are compiled from 2 independent experiments (n=9 mice per group). (C) AF488-Ovalbumin uptake in live sHz experienced DC2.4 cells. gMFI of AF488 is indicative of antigen uptake and measured by flow cytometry. (D) LNP mRNA uptake in live sHz experienced DC2.4 cells. gMFI of GFP is indicative of LNP mRNA uptake and translation of GFP mRNA. Data is compiled from 3 independent experiments (n=4-5 replicates per group). Dashed line is indicative of baseline gMFI of untreated cells.

nHz has been implicated in compromised antigen uptake by DCs^53,54^, but the impact of sHZ is unknown. To address this, we cultured DC2.4 cells with AF488 labeled whole ovalbumin protein and demonstrated a dose dependent inhibitory effect of sHz on ovalbumin uptake (**Fig. 6C**). Importantly, this effect was conserved in human monocyte derived DCs (**Extended Data Fig. 9**). To gain mechanistic insight into why LNP mRNA vaccination can overcome Hz induced immune suppression, we encapsulated a GFP encoding mRNA in our LNP formulation and measured LNP uptake in sHz exposed DC2.4 cells. We found equivalent expression of GFP in the DC2.4 cells regardless of sHz exposure (**Fig. 6D**). Thus, sHz does not impede LNP mRNA vaccine uptake, providing mechanistic insight into the ability of this vaccine to prime a response in malaria experienced hosts.

## DISCUSSION

Malaria is a devastating disease that kills more than 500,000 people annually^55^. A vaccine capable of taming this global health crisis remains frustratingly elusive. While RAS has been termed the “Gold Standard” of anti-malarial vaccines, its efficacy plummets in malaria-experienced individuals for reasons which are unclear^19,56^. We modeled impaired RAS efficacy in malaria experienced mice using *Py* or *Pcc* blood-stage infection prior to *Pb-*Ova RAS immunization, observing a marked reduction in antigen-specific CD8+ T cell responses in *Plasmodium* experienced mice. Due to the established importance of CD8+ T_EM_ and T_RM_ in controlling liver-stage malaria,^10,12^ we posit that defective CD8+ T cell responses are likely to blame for reduced RAS efficacy in the face of prior *Py* exposure.

Given that the impaired T cell phenotype was 1) observed with multiple strains of *Plasmodium* 2) durable— persisting for essentially the life of the animal, and 3) inducible by administering the contents of fixed iRBCs, we mechanistically focused our efforts to the long-lived and conserved biocrystal Hz. Hz has been reported to have a potential broad spectrum of biological activities including activation of the inflammasome pathway^57^, oxidative stress induction^58^, and activation of endothelial cells^59^. Several studies have indirectly assessed the contribution of Hz to immune impairment including its potential effects on DC function^60–62^. These studies have been complicated to interpret due to variations in the method of hemozoin preparation. Studies in this vein have utilized either injection or in vitro application of infected red cells^54,63,64^ or native or purified forms of Hz isolated from iRBCs, which often have parasite derived nucleic acids, lipids and proteins bound^65,66^ For example, one study identified Hz as a stimulator of TLR9, though upon treatment of the purified Hz with DNAse to eliminate contaminating parasite nucleotides, the TLR9 stimulation was abolished^67^. Moreover, native Hz complexed to parasite DNA has potent innate immune stimulating properties through NLRP3^28^ and can amplify innate immunity to liver-stage malaria for up to a month after inoculation^68^. However, our results showing that the CD8+ T cell defect was independent from the NLRP3 inflammasome suggests that Hz itself may have its own immunomodulatory properties, particularly since the defect is observed long after parasite clearance and with sHz. The consequences of Hz exposure long after parasite clearance remain a crucial topic in malaria biology.

We observed impaired CD8+ T cell responses and compromised RAS efficacy in mice treated with synthetic Hz, phenocopying our findings in *Plasmodium* experienced mice. While further studies are required to elucidate how exactly Hz is interacting with immune cells during critical steps in T cell activation, these initial studies point to Hz compromising DC functions, including antigen uptake. The generation of RAS-specific CD8+ T cell responses is dependent upon the acquisition of sporozoite antigens from dying hepatocytes by CD11c DCs^52^. Therefore, the persistence of Hz in the liver could be a driving force behind impaired immunity specifically observed in RAS vaccination in *Plasmodium* experienced individuals. Whether blood-stage malaria induces additional, non-Hz dependent long-lived immune alterations that compromise other vaccine responses remains an open question.

There are no therapeutic strategies shown to clear the insoluble biocrystal Hz from *Plasmodium* experienced hosts. Therefore, the best current option to rectify impaired immunity to liver-stage *Plasmodium* infection is to design vaccines that robustly function in Hz-laden hosts. DC peptide immunization was capable of eliciting robust CD8+ T cell responses in *Py* experienced mice, indicating that alternative vaccine strategies are possible. However, mRNA-based vaccines are likely to be a more clinically relevant strategy. We found mRNA vaccination led to equivalent vaccine specific CD8+ T cell responses between naïve and *Py* experienced mice. This finding supports the idea that clinically translatable alternative vaccine strategies could rectify Hz mediated suppression of CD8+ T cell responses to liver-stage malaria. A prior study utilized a whole protein (RPL6) encoding mRNA vaccine packaged in lipoplexes and similarly found that vaccination of *Py* experienced mice could yield protection from a *Pb* challenge^24^. Our study is consistent with this finding, though distinct in that our mRNA vaccine construct is designed to optimize MHC-I processing of multiple CD8+ T cell epitopes via ubiquitin degradation, as well as altered encapsulation chemistry (LNP vs. Lipoplex). Our study determined that while Hz impaired whole protein antigen uptake by DCs, our LNP formulation could bypass this impairment providing insight into why mRNA vaccination can overcome *Plasmodium* induced vaccine failure. Sophisticated studies in Rhesus macaques demonstrated that APCs are the primary cell of entry of LNP mRNA vaccines in vivo^69^. Future studies will be required to determine whether the mechanisms by which LNP vs. lipoplexes overcome Hz induced impairment of vaccination are the same. We have expanded upon the Ganley finding by utilizing the combined vaccination strategy of RAS and mRNA in a prime-pull approach provided enhanced protection, likely via enhanced liver T_RM_ formation even in *Plasmodium* experienced mice. Being poised at the site of infection, liver T_RM_ offer protection from liver-stage *Plasmodium*. This, in conjunction with the fact that RAS and LNP mRNA vaccines are fully translatable, make this vaccine strategy a particularly attractive one.

Ultimately, these studies identify Hz as a mechanistic driver of impaired RAS vaccine efficacy in malaria exposed individuals through the impairment of effector and memory CD8+ T cell responses. We showed that a combination mRNA+RAS vaccination regimen can produce robust T cell immunity and protection from malaria in murine hosts, regardless of exposure to Hz or prior malaria episodes. This finding offers hope that more effective malaria vaccines capable of circumventing Hz induced defects may be possible.

## METHODS

### Mice

C57BL/6J, OT-I, B6 Thy1.1 and NLRP3-/- mice were purchased from Jackson Labs and maintained at the University of Iowa husbandry facilities in accordance with approved IACUC protocols (#0051102). OT-I Thy1.1 mice were generated by crossing OT-I Thy1.2/1.2 mice with B6 Thy1.1/1.1 mice.

### Parasites

Sporozoites were harvested from infected *A. stephensi* mosquitoes generated in the Harty Lab insectary at the University of Iowa (*P. berghei* ANKA Ova-mCherry^30^, *P. yoelii* 17XNL) or procured from the University of Georgia SPOROcore (*P. berghei*-LUC). *P. berghei* ANKA Ova-mCherry sporozoites expresses OVA which is C-terminally fused to mCherry under the control of the 5′-UTR of hsp70 and the 3′-UTR of dhfr/ts. *P. berghei* sporozoite challenges used 1×10^4^ sporozoites, *P. yoelii* sporozoite challenges used 5×10^2^ sporozoites.

*P. yoelii* 17XNL was obtained from the NYU insectary and maintained as cryopreserved infected red blood cell stocks produced from a C57BL/6 mouse infected from sporozoites ^29^. 1×10^6^ cryopreserved iRBCs were used for blood-stage challenges via retroorbital intravenous injection. For all experiments, challenge was confirmed to result in parasitemia by collecting peripheral blood between days 7 and 15 post infection.

*P. c. chabaudi* AS-mCherry parasites were obtained from Patrick Duffy (NIAID) as cryopreserved stocks of infected red blood cells. Female BALB/c mice were used to revive cryopreserved iRBCs and produce fresh iRBCs for experiments. 1×10^6^ *P. c. chabaudi* AS-mCherry-infected iRBCs were used for challenge by intravenous injection at the retroorbital sinus. Chronic *P. c. chabaudi* AS-mCherry infection was cured by administering 600 μg of chloroquine intraperitoneally once daily for 10 days beginning at 15 DPI.

### Measurement of parasitemia

Breakthrough parasitemia in early blood-stage was assessed at day 5 post sporozoite challenge. Peripheral blood parasitemia was quantified by staining glutaraldehyde-fixed blood harvested via tail tip with Hoechst, anti-CD45 and anti-Ter-119, as previously described^52,70^. Infected RBCs were defined as Hoechst-positive/ CD45-negative/ Ter-119-positive cells.

### Radiation-attenuated sporozoite (RAS) immunization

*P. berghei* sporozoites were harvested from infected *A. stephensi* mosquito salivary glands and irradiated with 200 Gy (20,000 rads) to produce radiation attenuated sporozoites (*Pb-* RAS) as described^52^. Immunizations were performed by I.V. injection at the retro-orbital sinus with 1×10^4^ *Pb-*RAS.

### Liver parasite burden by qPCR

Liver parasite burden was determined as previously described^71,72^. Livers were extracted at 40-44 hours post sporozoite challenge, minced by GentleMACS processing (Miltenyi) and then digested in liver digest medium (ThermoFisher 17703034). Homogenized livers were pushed through 100 micron filters and hepatocytes collected by pelleting via centrifugation at 500 RPM. Hepatocyte pellets were resuspended in Trizol and snap frozen in liquid nitrogen. Total RNA was extracted following DNase digestion/cleanup with RNA clean and concentrator kit (Zymo). *Plasmodium* 18S rRNA was quantified by qPCR analysis on 2 microgram RNA samples using the Fast Virus One-step qPCR kit (Applied Biosystems). *Plasmodium* 18S rRNA values were normalized for input RNA with a GAPDH control.

### Liver parasite burden by IVIS

Liver-stage parasite burden was assessed by intravital luminescence imaging (IVIS) 40- 44 hours after challenge with luciferase-expressing sporozoites. Mice abdomens were shaved 1 day prior to IVIS imaging. To enable imaging, mice were injected intraperitoneally with inVivoglo D-luciferin (ProMega) and anesthetized with 2% isoflurane. Luminescence in equal-sized areas of interest were quantified using Xenogen Living Image software (Caliper Life Sciences).

### Transfer of OT-I transgenic CD8+ T cells

OT-I cells were harvested from naïve female donors by drawing peripheral blood. Vitalyse (CMDG) was used to remove red blood cells, and OT-I abundance determined by flow cytometry, staining for CD8-positive/CD90.1-positive cells. 10^4^ OT-I cells were transferred into recipients by IV injection at the retro-orbital sinus 1 day prior to immunization.

### Analysis of effector phase CD8+ T cell responses

Mice received naive OT-I cells as indicated, prior to immunization with 1×10^4^ *Pb-*RAS, expressing Ova as indicated. Peripheral blood was collected at day 6 or day 7, and stained with the following panel: CD8-BV785, CD11a-BV510/AF488, Thy1.1-PerCP Cy5.5/eFluor710, CD4- FITC, and APC-conjugated RPL6 tetramer.

### Analysis of memory phase CD8+ T cell responses

Thirty days post vaccination, mice were euthanized. Spleens were collected in RPMI and mashed over a 70 micron mesh filter to form a single cell suspension. Red blood cells were lysed using VitaLyse per manufacturer’s instructions. Cells were stained with the following panel to evaluate circulating memory T cell responses: CD8-AF700, CD11a-AF488, CD90.1- PerCP-Cy5.5, CD62L-PacBlue, and CD27-BV785. Memory OT-I were identified as CD8+, CD11ahi, and CD90.1+. Livers were collected into liver digest medium (ThermoFisher) and minced by GentleMacs (Miltenyi) processing. Homogenized livers were pushed through 100 micron filters and spun through a 35% percoll gradient. Red blood cells were lysed using VitaLyse per manufacturer’s instructions. Cells were stained with the following panel to evaluate liver T_RM_ responses: CD8-BV785, CD69-PE dazzle, CD90.1-AF700, CXCR6-AF647, CD62L-PacBlue, CD11a-AF488, and an efluor780 viability dye. OT-I T_RM_ were defined as: Live, CD8+, CD90.1+, CD69+, CXCR6+. To evaluate functionality, lymphocytes isolated from the livers were stimulated with 200 nM ova peptide for 6 hours in the presence of brefeldin A. Cells were surface stained with anti- CD8-BV785, CD11a-BV510, CD69-PE, CD62L-PE Dazzle, CXCR3- BV421, Thy1.1-FITC, and an efluor780 viability dye. Cells were fixed and stained with anti-IFNγ-APC prior to analysis. Cells were analyzed using a BD Fortessa flow cytometer.

### DC-Ova prime^73^

C57BL/6J mice were seeded with 5×10^6^ Flt3 ligand expressing B16 melanoma cells. 14 days post-seeding, mice were injected with 1 μg LPS IV. 24 hours later, splenocytes were harvested and CD11c+ DCs were enriched using a Miltenyi negative selection kit. Cells were pulsed with 8-mer Ova peptide and residual peptide was washed away. 500,000 peptide pulsed DCs were transferred intravenously to Py experienced or naïve mice.

### Histology

At various time points post *Plasmodium* exposure, mice were perfused with 20 ml of phosphate buffered saline, followed by 10 ml of 4% paraformaldehyde. Livers, spleens and femurs were collected into 4% paraformaldehyde and fixed overnight. Tissues were embedded in paraffin blocks for sectioning. 5 μm sections were mounted, processed and haematoxylin and eosin (H&E) stained and imaged on a slide scanning microscope.

### Hemozoin quantification

Hemozoin was isolated and quantified as previously described^74^. Briefly, tissue was collected into 5 ml of sterile water and homogenized. The homogenate was pelleted and washed with 2% SDS, 100 mM sodium bicarbonate and 2% SDS. The pellet was treated with proteinase K overnight at 60 degrees C. Following proteinase K digestion, the pellet was washed with water and decrystalized with 2% SDS, 20 mM NaOH for 1 hour at room temperature. Serial dilutions of the extracted hemozoin were made and absorbance 400 nm was measured with a spectrophotometer using a 1 cm cuvette. Hemozoin was quantified using a molar extinction coefficient of 10^5^.

### Beta-hematin hemozoin transfer

Beta-hematin (InvivoGen, sHZ) was prepared at 5 mg/mL in PBS and sonicated immediately prior to injection. Mice were given 1 mg doses of hemozoin via intravenous injection at the retro-orbital sinus, with 5 doses over the course of 1 week.

### Transfer of fixed parasitized RBCs

Female C57BL/6 mice were infected with 1×10^6^ *P. berghei*-infected RBCs. Peripheral blood from infected or influenza PR8 immune control donors was collected, fixed with 0.635% glutaraldehyde, washed 3x with PBS, and then 1×10^6^ fixed iRBCs were transferred into naive recipient mice by intravenous injection. Lack of viability in the transferred fixed iRBCs was confirmed by monitoring recipient mice for parasitemia.

### Antigen uptake assay

Monocytes were enriched from PBMCs using a Miltenyi human pan monocyte isolation kit and plated in differentiation medium (RPMI supplemented with 10% FBS, 1% pen/strep, 1% sodium pyruvate, 1% HEPES, 1% non-essential amino acids, 50 ng/ml GM-CSF, and 50 ng/ml IL-4) for 6 days. Cells were collected and phenotype was confirmed by flow cytometry for CD14, CD16, CD45, and CD11c expression. Human monocyte derived DCs or mouse DC2.4^75^ cells were plated and activated with 1 μg/ml of LPS for 24 hours in the presence or absence of sHz. Residual sHz was washed from the cells with PBS. Ovalbumin-AF488 or LNPs containing GFP encoding mRNA was pre-incubated with a 1:1 volume ratio of mouse serum at 37°C for 15 minutes. 4 μg/ml of Ovalbumin-AF488 or 1 μg/ml of LNPs in complete RPMI were added were added to the cells After 1.5 hours (Ovalbumin-AF488) or 24 hours (GFP LNP), cells were harvested, stained for viability (efluor780) and analyzed by flow cytometry.

### mRNA vaccine preparation

A minigene insert was cloned into the pMRNAxp expression cassette driven by a type II T7 promoter element. The expression cassette encoded an A76 mutant ubiquitin (to drive more efficient proteasomal processing)^76^ followed by a flexible linker and a string of H-2^b^ and H-2^d^ restricted *P. berghei* optimal 8/9-mer CD8+ T cell epitopes (S20, Trap130, CS252, RPL6, and Ova). Each epitope was flanked by an AAY optimal proteasomal cleavage residue to aid in efficient proteasomal processing. In vitro mRNA expression was performed using an NEB-ARCA T7 kit supplemented with 50% ψUTP and 5mCTP. RNA was purified using an RNeasy mini kit (Qiagen) and RNA integrity was validated by RNA bleach gel. mRNA was packaged into lipid nanoparticles (LNP) using a Precigenome Nanogenerator Flex microfluidics device at a flow rate of 3:1 (aqueous:organic). The organic phase was prepared by combining a neutral lipid mixture (LipidFlex) with cationic lipid SM-102 at a molar ratio of 6:4, respectively. The aqueous phase was prepared by diluting the mRNA in 50 mM sodium acetate (pH 5.0). The N/P ratio was fixed at 6 for all LNP preparations. The LNP size was measured at 90-125 nm using a Zetasizer Nano ZS. The encapsulation efficiency and encapsulated mRNA concentration was confirmed using a Ribogreen encapsulation assay^77^. Mice were vaccinated intravenously with 5 μg of encapsulated mRNA in the retro-orbital sinus at the time of RAS immunization.

### Statistics

Statistical analyses were performed in Prism (Graphpad). For comparisons between 3-9 groups, an ANOVA statistical test with post hoc analysis (Tukey’s multiple comparison test) was utilized to determine statistical significance. For comparisons between 2 groups, a Mann-Whitney test was utilized to determine statistical significance. Statistical means of each group are represented as the line for each group. Experiments were repeated at least 2 times with an n of greater than or equal to 4 animals per group.

## Abbreviations

(T_RM_): Tissue resident memory T cell
(T_circM_): circulating memory T cell
(RAS): radiation attenuated sporozoite
(Hz): hemozoin
(DPI): days post infection
(*Pb*): *P. berghei*
(*Py*): *P. yoelii*
(*Pcc*): *P. chabaudi chabaudi*
(CQ): chloroquine

## Acknowledgements

We thank members of the Harty Lab, Noah Butler (University of Iowa), Vladimir Badovinac (University of Iowa) and Sam Kurup (University of Georgia) for fruitful discussions. We thank Ivan Badovinac, Cassie Sievers, Jack Harty (no relation), and Zachary Darr for maintaining laboratory solutions and equipment. Patrick Duffy (NIAID) generously shared *P. c. chabaudi* parasites. IVIS experiments were performed at the University of Iowa Small Animal Imaging Core. The University of Georgia SporoCore provided *P. berghei*-LUC sporozoites. The University of Iowa Free Radical and Radiation Biology Ionizing Radiation Services Core performed sporozoite radiations. Human MoDCs were differentiated from PBMCs obtain from the McGowin Blood Center by the University of Iowa Human Immunology Core. Funding was provided by NIH Grants to JTH (AI42767, AI100527, AI114543, AI167847, and AI185725), NIH (R21 EY034198-01; R21ES032937 - 01A1; R21 DE031042-01A1; 2P30CA086862), Lyle and Bighley Chair of Pharmaceutical Sciences, National Science Foundation (2242763) to AKS, Environmental Health Sciences Research Center (P30ES005605) to AKS and RH. MH was supported by NIH fellowship grants T32AI007260, 1F32AI174382, and 1 K99 AI190129-01 and the Burroughs Wellcome Fund grant number 1051302. LLD was supported by NIH fellowship grants T32AI007260 and 1F32AI167088. SA and MRM were supported by the University of Iowa MSTP Training Grant T32 GM139776. MRM was supported by the University of Iowa Graduate College Post-Comprehensive Research Fellowship.

## Author contributions

M.H.- Conceptualization, data curation, formal analysis, funding acquisition, investigation, methodology, validation, visualization, writing (original draft), and writing (review and editing). L.L.D.- Conceptualization, data curation, formal analysis, funding acquisition, investigation, methodology, validation, writing (review and editing). L.P.E.- Investigation and writing (review and editing). L.S.H.- Investigation, methodology, and writing (review and editing). R.H.- Methodology, resources, and writing (review and editing). M.R.M.- Investigation and writing (review and editing). S.A.- Investigation and writing (review and editing). A.K.S.- Funding acquisition, project administration, resources, supervision, and writing (review and editing). J.T.H.- Conceptualization, funding acquisition, project administration, resources, supervision, writing (original draft), and writing (review and editing).

## Extended Data Figure Legends

**Extended Data Fig. 1:** Gating strategy for CD8+ T cells. 7 (effector) or 30 (memory) days post immunization, PBL or splenocytes, respectively were collected and stained using anti-CD8 AF700, anti-CD11a AF488, anti-Thy1.1 Percp Cy5.5, and RPL6 specific MHC-I tetramer APC. Representative liver T_RM_ gating strategy. 30 days post immunization, livers were harvested from mice, processed, stained, and analyzed by flow cytometry. Cells were gated by FSC/SSC to identify lymphocytes, followed by a live cell gate. Cells were then gated on CD8+ and CD90.1+ to identify transgenic OT-I cells (Fig. 1). Liver T_RM_ were identified as CD69+ CXCR6+ (Fig. 4).

**Extended Data Fig. 2:** Parasite load during RAS immunization is equivalent between naïve and *P. yoelii* experienced mice. 18 hours post RAS immunization, livers were harvested and relative antigen load was measured using qRT-PCR. Data are representative of 2 independent experiments (n=3 mice per group).

**Extended Data Fig. 3:** Even with multiple RAS immunizations, *Py* exposed mice never fully catch up to age-matched controls. Mice were immunized with 10^4^ *Pb*-Ova RAS, allowed to rest 30 days and immunized a second time. 30 days after the last immunization, mice were challenged with 10^4^ virulent *Pb* sporozoites. Relative liver parasite burden 40-44 hours after virulent *Pb* challenge (parasite 18s RNA/host GAPDH) was measured by qRT-PCR. Data are pooled from 3 independent experiments (n=18-20 mice per group).

**Extended Data Fig. 4:** Live parasites are not required for observed T cell defect. C57BL/6 mice were injected with 10^6^ glutaraldehyde fixed iRBCs from *Pb* patent mice or an equivalent amount of RBCs from an influenza A virus immune mouse. 30 days after exposure, 10,000 OT-I cells were transferred to each mouse and mice were immunized with 10^4^ *Pb* Ova RAS. 7 days post immunization, effector OT-I cells were quantified in PBL. Data are representative of 2 independent experiments (n=4-5 mice per group).

**Extended Data Fig. 5:** sHz quenches luciferase signal. C57BL/6 mice were injected IV with 1 mg of sHz five times over the course of 1 week, totaling 5 mg of sHz administered per mouse. 60 days after the last sHz dose, mice were challenged with 10^4^ virulent *P. berghei*-Luc. In vivo luciferase expression was measured at 42 hours post infection. Data are from 2 independent experiments (n=11-13 mice per group).

**Extended Data Fig. 6:** Anti-ovalbumin IgG responses. 25 days following immunization, serum was collected from mice. Serial dilutions of serum were utilized in a whole ovalbumin IgG ELISA. Area under the curve measurements for A450 are displayed. Data are from 2 independent experiments (n=3-5 mice per group).

**Extended Data Fig. 7:** Prime-pull vaccine strategy yields enhanced liver T_RM_ formation in sHz exposed mice. C57BL/6 mice were treated with 5 mg of sHz 1 month prior to immunization. 10,000 OT-I cells were transferred to each mouse. Mice were vaccinated with either 10^4^ *Pb* Ova RAS, 5 μg of Ub *Pb* Ova vaccine, or both. 30 days post immunization, spleens (A) and livers (B) were harvested. Memory CD8+ T cell responses were quantified by flow cytometry. Data is pooled from 2 independent experiments (n=6 per group).

**Extended Data Fig. 8:** Observed T cell defect is not dependent upon NLRP3 inflammasome. WT or NLRP3-/- mice were exposed to *Py* iRBCs. 30 days following infection, 10,000 OT-I cells were transferred to each mouse and mice were immunized with 10^4^ P. berghei RAS. 7 days post immunization, effector OT-I cells were quantified in PBL. Data are pooled from 3 independent experiments (n=7-11 mice per group).

**Extended Data Fig. 9:** sHz impairs human monocyte derived DC antigen uptake. AF488- ovalbumin uptake by human monocyte derived DCs. (n=3 replicates per group).

## Data Availability Statement

All data supporting the findings of this study are available within the paper and its Supplementary Information files. Source data are provided with this paper. Additional data that support the conclusions of this study are available from the corresponding author upon reasonable request.

