## Supplemental data for "mRNA vaccination overcomes hemozoin-mediated impairment of whole parasite vaccine efficacy for malaria"

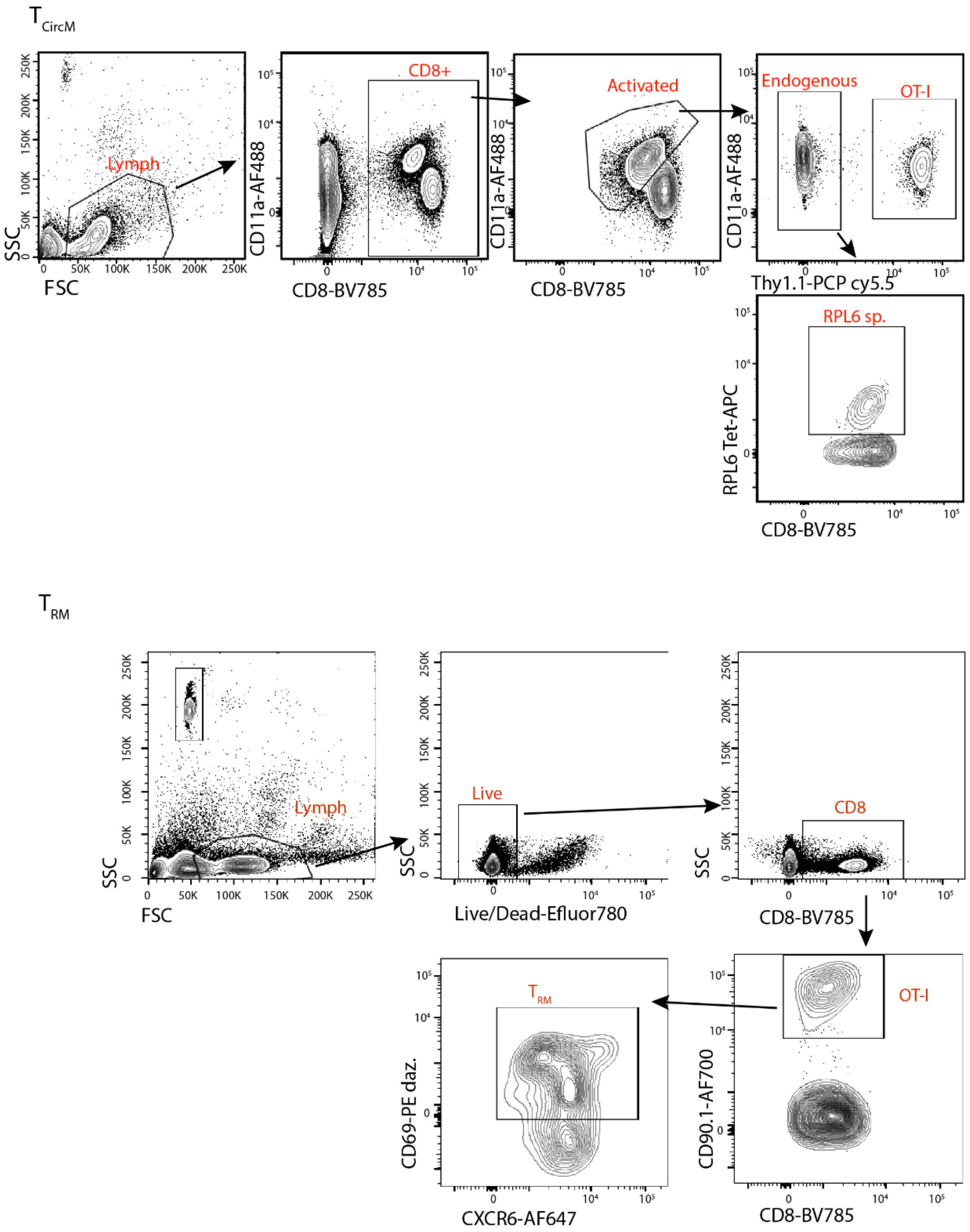


**Extended Data Fig. 1:** Gating strategy for CD8+ T cells. 7 (effector) or 30 (memory) days post immunization, PBL or splenocytes, respectively were collected and stained using anti-CD8 AF700, anti-CD11a AF488, anti-Thy1.1 Percp Cy5.5, and RPL6 specific MHC-I tetramer APC. Representative liver $T_{RM}$ gating strategy. 30 days post immunization, livers were harvested from mice, processed, stained, and analyzed by flow cytometry. Cells were gated by FSC/SSC to identify lymphocytes, followed by a live cell gate. Cells were then gated on CD8+ and CD90.1+ to identify transgenic OT-I cells (Fig. 1). Liver $T_{RM}$ were identified as CD69+ CXCR6+ (Fig. 4).

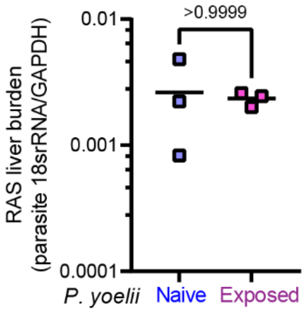

**Extended Data Fig. 2:** Parasite load during RAS immunization is equivalent between naïve and *P. yoelii* experienced mice. 18 hours post RAS immunization, livers were harvested and relative antigen load was measured using qRT-PCR. Data are representative of 2 independent experiments (n=3 mice per group).

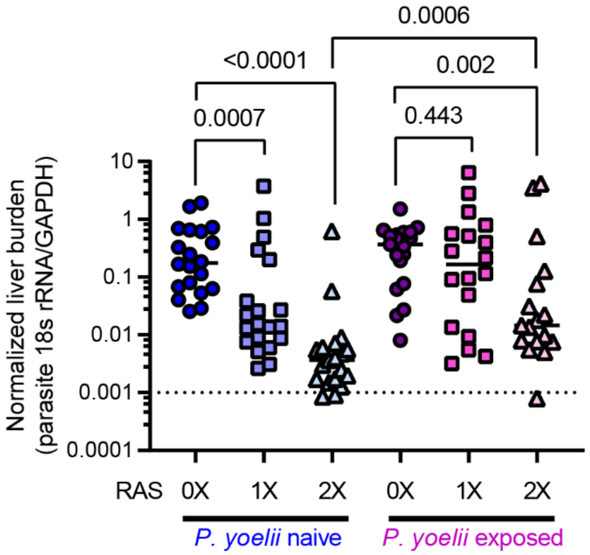


**Extended Data Fig. 3:** Even with multiple RAS immunizations, *Py* exposed mice never fully catch up to age-matched controls. Mice were immunized with $10^4$ *Pb*-Ova RAS, allowed to rest 30 days and immunized a second time. 30 days after the last immunization, mice were challenged with $10^4$ virulent *Pb* sporozoites. Relative liver parasite burden 40-44 hours after virulent *Pb* challenge (parasite 18s RNA/host GAPDH) was measured by qRT-PCR. Data are pooled from 3 independent experiments (n=18-20 mice per group).

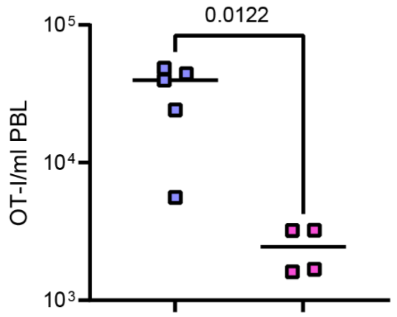


**Extended Data Fig. 4:** Live parasites are not required for observed T cell defect. C57BL/6 mice were injected with $10^6$ glutaraldehyde fixed iRBCs from *Pb* patent mice or an equivalent amount of RBCs from an influenza A virus immune mouse. 30 days after exposure, 10,000 OT-I cells were transferred to each mouse and mice were immunized with $10^4$ *Pb* Ova RAS. 7 days post immunization, effector OT-I cells were quantified in PBL. Data are representative of 2 independent experiments (n=4-5 mice per group).

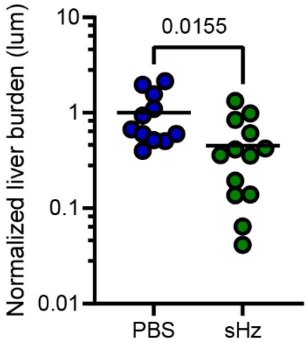

**Extended Data Fig. 5:** sHz quenches luciferase signal. C57BL/6 mice were injected IV with 1 mg of sHz five times over the course of 1 week, totaling 5 mg of sHz administered per mouse. 60 days after the last sHz dose, mice were challenged with $10^4$ virulent *P. berghei*-Luc. In vivo luciferase expression was measured at 42 hours post infection. Data are from 2 independent experiments (n=11-13 mice per group).

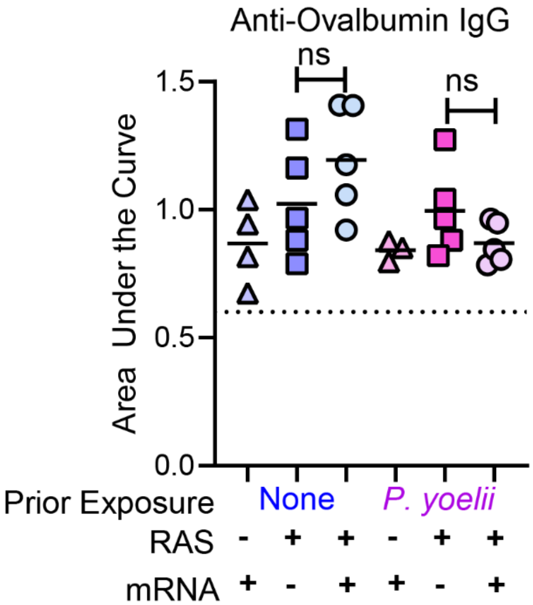


**Extended Data Fig. 6**: Anti-ovalbumin IgG responses. 25 days following immunization, serum was collected from mice. Serial dilutions of serum were utilized in a whole ovalbumin IgG ELISA. Area under the curve measurements for A450 are displayed. Data are from 2 independent experiments (n=3-5 mice per group).

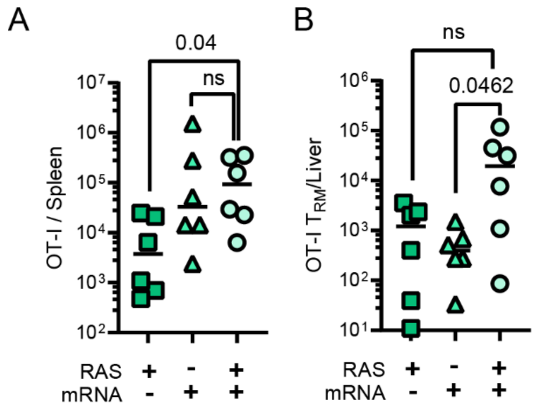


**Extended Data Fig. 7**: Prime-pull vaccine strategy yields enhanced liver $T_{RM}$ formation in sHz exposed mice. C57BL/6 mice were treated with 5 mg of sHz 1 month prior to immunization. 10,000 OT-I cells were transferred to each mouse. Mice were vaccinated with either $10^4$ *Pb* Ova RAS, 5 µg of Ub *Pb* Ova vaccine, or both. 30 days post immunization, spleens (A) and livers (B) were harvested. Memory CD8+ T cell responses were quantified by flow cytometry. Data is pooled from 2 independent experiments (n=6 per group).

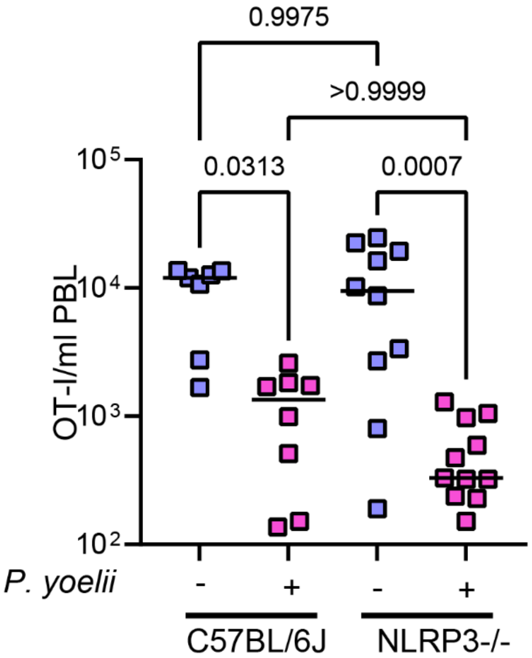


**Extended Data Fig. 8:** Observed T cell defect is not dependent upon NLRP3 inflammasome. WT or NLRP3-/- mice were exposed to *Py* iRBCs. 30 days following infection, 10,000 OT-I cells were transferred to each mouse and mice were immunized with $10^4$ P. berghei RAS. 7 days post immunization, effector OT-I cells were quantified in PBL. Data are pooled from 3 independent experiments (n=7-11 mice per group).

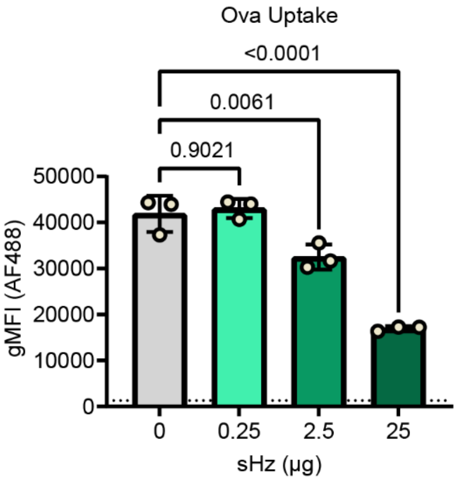


**Extended Data Fig. 9:** sHz impairs human monocyte derived DC antigen uptake. AF488-ovalbumin uptake by human monocyte derived DCs. (n=3 replicates per group).
